# RudLOV—a new optically synchronized cargo transport method reveals unexpected effect of dynasore

**DOI:** 10.1101/2023.11.04.565648

**Authors:** Tatsuya Tago, Takumi Ogawa, Yumi Goto, Kiminori Toyooka, Takuro Tojima, Akihiko Nakano, Takunori Satoh, Akiko K. Satoh

**Affiliations:** Division of Life Science, Graduate School of Integral Arts and Science, Hiroshima University, Higashi-Hiroshima, Hiroshima 739-8521, Japan; Technology Platform Division, Mass Spectrometry and Microscopy Unit, RIKEN Center for Sustainable Resource Science, Yokohama, Kanagawa 230-0045, Japan; Live Cell Super-Resolution Imaging Research Team, RIKEN Center for Advanced Photonics, 2-1 Hirosawa, Wako, Saitama 351-0198, Japan

## Abstract

Live imaging of secretory cargoes is a powerful method for understanding the mechanisms of membrane trafficking. Inducing the synchronous release of cargoes from an organelle is a key for enhancing microscopic observation. We developed an optical cargo-releasing method named as retention using dark state of LOV2 (RudLOV), which enables exceptional spatial, temporal, and quantity control during cargo release. A limited amount of cargo-release using RudLOV successfully visualized cargo cisternal-movement and cargo-specific exit sites on the Golgi/*trans*-Golgi network. Moreover, by controlling the timing of cargo-release using RudLOV, we revealed the canonical and non-canonical effects of the well-known dynamin inhibitor dynasore, which inhibits early-Golgi but not late-Golgi transport and exit from the *trans*-Golgi network where dynamin-2 is active. Accumulation of COPI vesicles at the *cis*-side of the Golgi stacks in dynasore-treated cells suggests that dynasore targets COPI-uncoating/tethering/fusion machinery in the early-Golgi cisternae or endoplasmic reticulum but not in the late-Golgi cisternae. These results provide insight into the cisternal maturation of Golgi stacks.

## Introduction

Most membrane and secretory proteins are synthesized in the endoplasmic reticulum (ER), transported to the Golgi apparatus, and then travel to their destinations. Several components involved in these processes in various secretory pathways have been identified and characterized biochemically and structurally. Recent advances in optical and electron microscopy revealed new aspects of membrane interactions and cargo transfer (Kurokawa *et al*, 2014; Shomron *et al*, 2021; Solinger *et al*, 2020; Weigel *et al*, 2021). Tracking secretory cargoes is a key technique in microscopic analyses and requires pulse labeling or the synchronous release of cargo proteins for clear visualization (Farr *et al*, 2009; Kurokawa *et al*, 2019; Lippincott-Schwartz *et al*, 2000; Presley *et al*, 1997).

Accordingly, several regulatory secretory cargoes and cargo retention and release systems have been developed (Boncompain *et al*, 2012; Bourke *et al*, 2021; Casler *et al*, 2020). The most widely used method is the retention using a selective hook (RUSH) system, which uses streptavidin, streptavidin-binding peptide (SBP), and biotin (Boncompain *et al*., 2012). Streptavidin tagged an organelle-specific retention signal traps SBP-tagged cargo proteins. Addition of biotin to the medium triggers the synchronous release of SBP-tagged cargo proteins from the organelle because biotin occupies the SBP-binding sites of streptavidin with high affinity. Cargo proteins fused to fluorescent proteins or tags allow for live imaging of cargoes during the journey from trapped organelle to the destination. However, the RUSH system is limited by its lack of spatial and quantity control for releasing cargos; supplying biotin to specific areas of cells is difficult, and stopping cargo release by washing out the biotin is nearly impossible because of the strong affinity of biotin for streptavidin, which results in prolonged cargo release. A more precise and controlled method of cargo release using 405-nm light illumination known as the zapalog-mediated ER trap was recently developed (Bourke *et al*., 2021). As illumination can be precisely controlled at the single-cell or subcellular domain levels using the same microscope used subsequent live imaging, cargo release can be triggered with exceptional spatial and temporal control. In fact, Bourke et al. visualized and compared the trafficking pathways of synaptic proteins in neurons after the release of these proteins from the ER located in the central (cell body) and local (dendritic) regions. One limitation of this method is that a continuous supply of relatively expensive zapalogs is required after transfection or expression of cargo proteins to trap the cargo within the ER. In addition, the limited permeability of the chemicals makes it difficult to apply this method to whole animals and tissues. Furthermore, 405 nm light used to degrade zapalogs can damage cells, although not to the same extent as the damage caused by UV light.

In the present study, we report a light-induced cargo release method named as retention using dark state of LOV2 (RudLOV). RudLOV is based on LOVTRAP, which is an optogenetic approach for reversible light-induced protein dissociation (Wang *et al*, 2016). RudLOV utilizes LOV2 as the hook in the ER. The cargoes fused with zdk1 selectively bind to the dark state of the LOV2 hook to be retained in the ER in the dark. Illumination at 443 nm triggers Zdk1-cargo release from the ER.

The ubiquitously expressed dynamin-2 plays an essential role in clathrin-mediated endocytosis, particularly in the scission of coated pits and vesicles from the plasma membrane (Ferguson & De Camilli, 2012). Several studies reported that dynamin-2 is also required for post-Golgi transport; however, its precise roles in the Golgi stacks remain unclear (Cao *et al*, 2005; Jones *et al*, 1998; Kessels *et al*, 2006; Kreitzer *et al*, 2000; Salvarezza *et al*, 2009). Indeed, dynamin-1/2 viable double KO cells show accumulation of clathrin coated pits on the plasma membrane but not on the TGN or Golgi stacks (Ferguson *et al*, 2009). Thus, we investigated the effect of the well-known dynamin inhibitor dynasore (Macia *et al*, 2006) on cargo transport using RudLOV. Precise temporal control of cargo transport enabled by RudLOV allowed us to dissect the steps of cargo transport within Golgi and *trans*-Golgi network (TGN), which revealed an unexpected effect of dynasore on the *cis*-side in addition to the *trans*-side of Golgi stacks.

## Results and discussion

### RudLOV enables highly controlled cargo-release with spatial and temporal precision

The most widely used method for the synchronous release of cargo proteins is the RUSH system, which is convenient because cargo release is achieved simply by adding biotin to the medium. However, we observed an unexpected variation in the response of the cells to biotin. HeLa cells showing high levels of cargo expression after transient transfection with RUSH constructs tended to exhibit a longer delay in inducing cargo export from the ER after biotin administration (Figure 1D, E). This lack of temporal control in the RUSH system coupled with the lack of spatial and quantity controls for cargo release prompted us to develop a new method for the synchronous release of cargo proteins.

**Figure 1.**
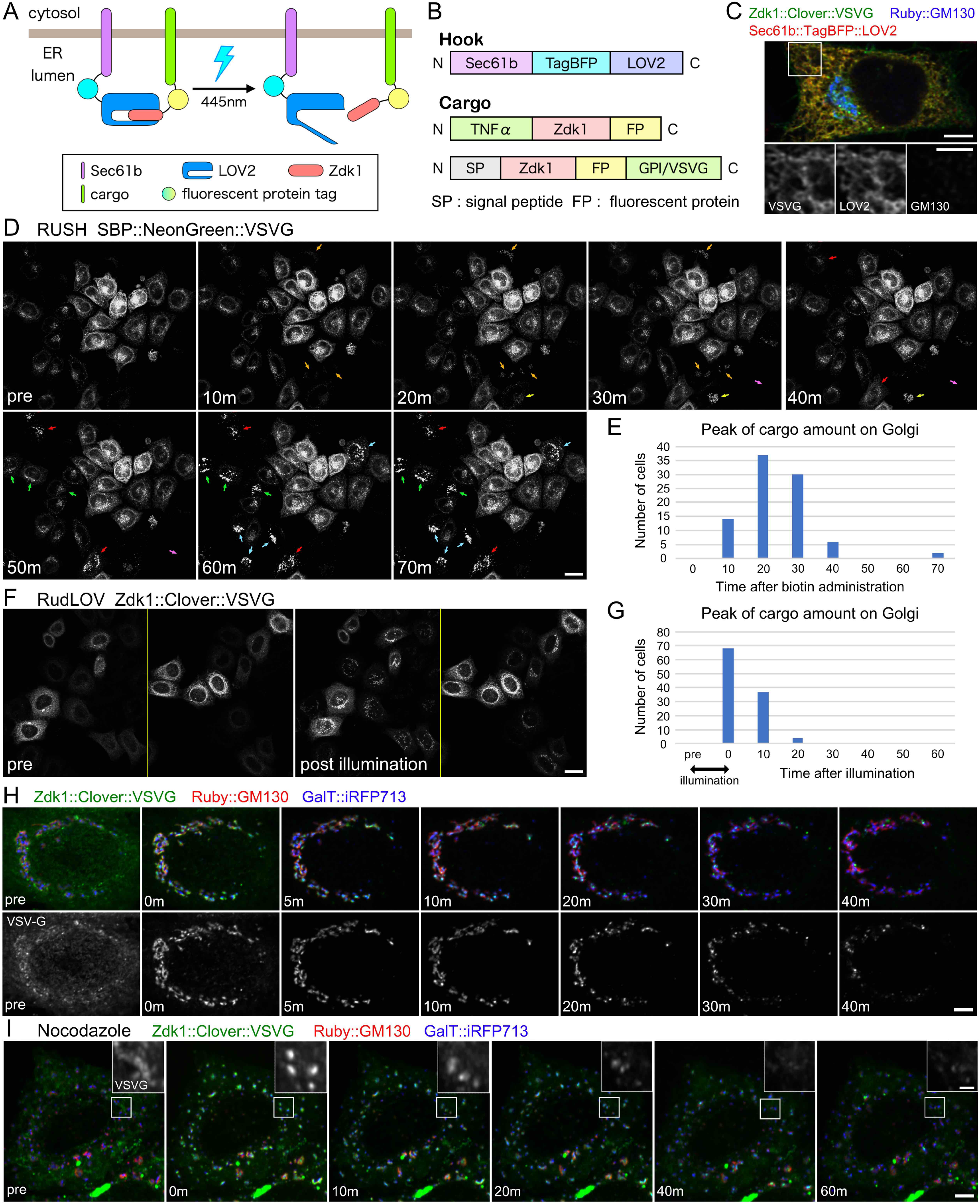
RudLOV enables highly synchronized cargo-release. (A, B) Schematic of the RudLOV method, and constructs of hook and cargoes. The N-terminal of LOV2 is fused with human Sec61b and fluorescent protein (TagBFP) to localize on the endoplasmic reticulum (ER). Cargoes and fluorescent proteins (Scarlet, Clover or NeonGreen) are fused at either the N- or C-terminus of Zdk1, which binds to LOV2 in the dark and detaches on illumination at 445 nm. (C) Localization of the hook Sec61b::tagBFP2::LOV2 (red) and cargo Zdk1::Clover::VSVG (green) in the dark condition. Golgi apparatus is marked by GM130 (blue). (D) SBP::NeonGreen::VSVG localizations before (left) and at 10, 20, 30, 40, 50, 60, and 70 min after addition of 100 μM of biotin in the RUSH system. Arrows indicate the cells with accumulated SBP::NeonGreen::VSVG on the Golgi apparatus. (E) Plot of cell counts showing peak amount of cargo (VSVG) on the Golgi apparatus at each time point after biotin administration in RUSH system. (F) Zdk1::Clover::VSVG localizations before (left) and after (right) illumination at 445 nm using the RudLOV system. (G) Plot of cell counts showing peak amount of cargo (VSVG) on the Golgi apparatus at each time point after illumination in RudLOV. (H, I) Localization of Zdk1::Clover::VSVG before (left) and at 0, 5, 10, 20, 30, and 40 min after illumination at 445 nm using the RudLOV system in untreated (H) and nocodazole-treated cells (I). Inset in I shows a magnified image of Zdk1::Clover::VSVG. Scale bars: 5 μm (C), 2 μm (inset of C), 20 μm (D, F), 5 μm (H, I), and 2 μm (inset of I).

The schematics of the RudLOV method and constructs are shown in Figure 1A and B. The LOV2 C-terminal helix Jα, which consists of the interface recognized by Zdk1, unwinds upon illumination. LOV2 is fused to the lumenal C-terminus of human Sec61b, which is a subunit of the translocon for ER-localization. Cargoes and fluorescent proteins were tagged with Zdk1 at either the N- or C-terminus. In the dark, the cargoes were retained in the ER by the LOV2 hook (Figure 1C). Upon illumination at 445 nm, the cargoes moved to the Golgi apparatus in all illuminated cells (Figure 1F). To avoid LOV2 activation during cargo detection, we used a 514 nm laser rather than a 488 nm laser to observe the Clover fluorescent proteins. To achieve seamless application of LOV2 activation and three-color imaging, we equipped an FV3000 confocal microscope with a quinta-band dichroic mirror for the 405/445/514/561/647 nm laser lines. This allowed for high-speed imaging without misalignment of the optical axis.

Three types of cargoes, glycosylphosphatidylinositol-anchored protein (GPI-AP), vesicular stomatitis virus protein G (VSVG), and tumor necrosis factor α (TNFα), were prepared (Figure 1B). After 10 min of illumination, all three cargoes accumulated in the Golgi apparatus, and most of them exited from Golgi by 40 min (Figure 1H and S1A, C). We also observed cargo movement in dispersed Golgi stacks after nocodazole treatment. All three types of cargoes accumulated in the Golgi apparatus after only 10 min of illumination, and most of them exited from Golgi by 40 min (Figure 1I and S1B, D), similar to the results in nocodazole-untreated cells (Figure 1H and S1A, C). This result is consistent with that of a previous study in which transport was observed using the RUSH system and showed that microtubules are not strictly required for secretion (Fourriere *et al*, 2016).

Next, we investigated the ability of RudLOV to activate localized cargo export. We found that RudLOV activated cargo export in just a single cell within a field (Figure 2A) or only within a single Golgi stack, although some leaks were observed in neighboring Golgi stacks (Figure 2B). Thus, RudLOV provides exceptional spatial and temporal secretion control. Notably, the RudLOV differed from the RUSH system in the manner of TNFα localization before cargo release (Figure 2C). TNFα is concentrated in the ER exit sites (ERES) in the RUSH system but diffused throughout the ER in RudLOV. This difference in the localization of ER-retained TNFα may reflect differences in the ER-localization signals of the hooks, which is the KDEL in RUSH and Sec61b in RudLOV. Sec61b but not KDEL may have sufficient activity to exclude TNFα from ERES.

**Figure 2.**
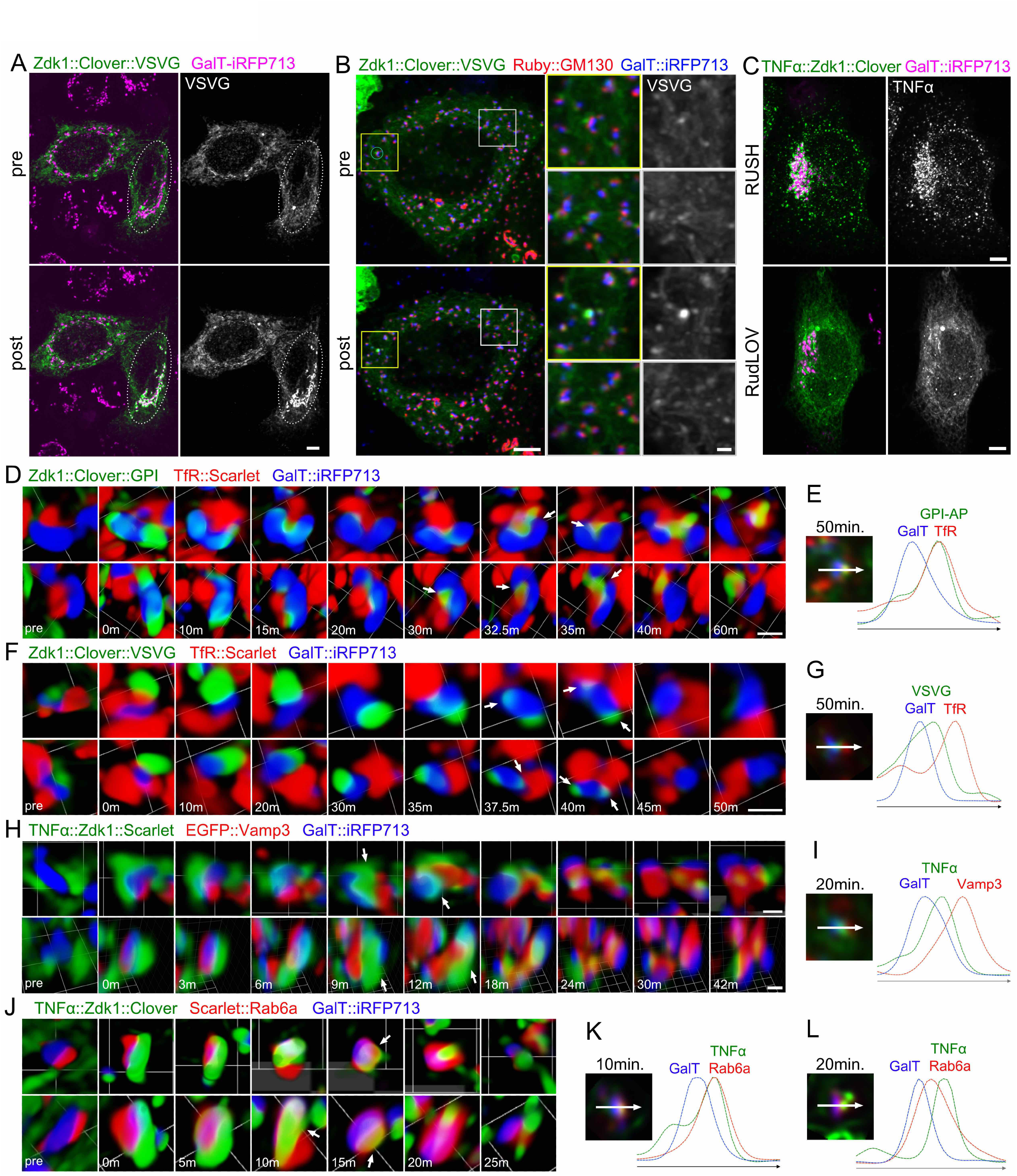
Localized activation of secretion using RudLOV. (A, B) Localization of the cargo Zdk1::Clover::VSVG approximately 10 min after illumination in untreated (A) and nocodazole-treated cells (B). The cargo Zdk1::Clover::VSVG is shown in green, and the *trans*-Golgi marker GalT::iRFP713 in magenta (A) or blue (B). The right cell but not the left cell is illuminated (A). A single Golgi stack located in the middle of the yellow inset is illuminated (B). (C) Localization of the cargoes TNFα::SBP::NeonGreen (upper) and TNFα::Zdk1::Clover (lower) before transport was triggered. Cargoes are presented in green and the *trans*-Golgi marker GalT::iRFP713 in magenta. (D–L) Localization of the cargo Zdk1::Clover::GPI (D, E), Zdk1::Clover::VSVG (F, G), TNFα::Zdk1::Scarlet (H, I) or TNFα::Zdk1::Clover (J–L) before (left) or after illumination in a single Golgi/RE unit in nocodazole-treated cells. One each of time lapse 3D volume containing a representative Golgi/RE unit is presented as pairs of time-series volumetrically rendered from two different view angles (D, F, H, J). The time after illumination is shown in the bottom-left corner. Plots show signal intensities from the image on the left Golgi/RE unit at indicated time points after illumination (E, G, I, K, L). Signal intensity was measured along the arrow (representing 1.5 μm). Arrows indicate Zdk1::Clover::GPI is on RE (D) and Zdk1::Clover::VSVG is not on RE (E). Arrows indicate TNFα::Zdk1::Scarlet on the membrane between *trans*-Golgi cisternae and RE (H) and TNFα::Zdk1::Clover is on Rab6-positive compartment (J). Cargoes are shown in green, a *trans*-Golgi marker GalT::iRFP713 in blue, and the RE marker TfR::Scarlet (D– G), RE marker GFP::Vamp3 (H, I), or early-TGN marker Scarlet::Rab6 (J–L) in red. Scale bars: 10 μm (A), 5 μm (B, C), 2 μm (inset in B) and 1 μm (D, F, H, J).

### TNFα and VSVG accumulate in the membrane between *trans*-cisternae and Golgi-associated recycling endosomes before exiting the Golgi stacks

We previously observed a relationship between the Golgi stacks and recycling endosomes (RE) in nocodazole-treated cells and described two types of REs: free RE and Golgi-associated RE (GA-RE). Most Golgi stacks are accompanied by GA-RE (Fujii *et al*, 2020a; Fujii *et al*, 2020b). Here, we refer to the Golgi stack/GA-RE complex as the Golgi/RE unit. We previously showed that GPI-AP is transported from the Golgi stack to the GA-RE before exiting the Golgi/RE unit, whereas VSVG did not. To understand the details of cargo transport within the Golgi/RE unit in nocodazole-treated HeLa cells, we visualized the movement of small amounts of cargo released under limited illumination. Prior to illumination, GPI-AP, VSVG, and TNFα were not detected around the Golgi stacks (Figure 2D, F, H, J left). After 5 min of illumination, GPI-AP was detected near the Golgi stack. Golgi stacks were accompanied by GA-RE during cargo secretion, and GPI-AP moved seamlessly from Galactosyltransferase (GalT) positive *trans*-Golgi cisternae to transferrin receptor (TfR)-marked GA-RE (Figure 2D, E). After 5 min of illumination, VSVG was also detected around the Golgi stack but did not enter the TfR-marked GA-RE even at 30 min after illumination (Figure 2F, G). VSVG remained near the *trans*-Golgi cisternae and GA-RE until around 40 min (Figure 2F). This result is consistent with those of a previous study in which the RUSH system was used (Fujii *et al*., 2020a). Interestingly, TNFα appeared to force GA-RE away to create more space for it to remain in the gap (Figure 2H arrow) between the *trans*-Golgi cisternae and GA-RE after its exit from the *trans*-Golgi cisternae (Figure 2H, I). To address the identity of the TNFα-positive membrane in the gap, we visualized the Rab6-positive compartment. Rab6 is previously described to locate on early-TGN in budding yeast (Tojima *et al*, 2019) and on the boundary between trans-cisternae and TGN in mammalian cells (Tie *et al*, 2018). TNFα seamlessly moved from GalT-positive *trans*-Golgi cisternae to the Rab6-positive early-TGN (Figure 2J, K), and then passed the early-TGN in approximately 20 min (Figure 2J, L). TNFα remained in an unidentified compartment attached to the Rab6-positive early-TGN until it exited the Golgi/RE unit. This unidentified compartment may be the late-TGN defined by clathrin and GGA in yeast (Tojima *et al*., 2019). Clathrin and GGA are also shown to be localized on the punctate structures within TGN but not adjacent to *trans*-cisternae in mammalian cells (Tie *et al*., 2018).These results indicate that the ability of RudLOV to precisely control the amount and timing of cargo release is useful for understanding cargo movement.

### Dynasore inhibits early-stage of intra-Golgi transport

Dynamin-2 is required for the formation of post-Golgi vesicles in the TGN (Cao *et al*., 2005; Jones *et al*., 1998; Kessels *et al*., 2006; Kreitzer *et al*., 2000; Salvarezza *et al*., 2009). A previous study showed that administration of dynasore to the cells inhibited cargo transport on or around the Golgi apparatus; however, a detailed analysis has not been performed (Weller *et al*, 2010). Hence, we used RudLOV to investigate the effects of dynasore on cargo transport. We observed cargo movement within the dispersed Golgi/RE unit in nocodazole-treated cells. Without dynasore, VSVG accumulated at the *cis*-side of the Golgi stacks immediately after illumination (defined as 0 min) but colocalized poorly with the *cis*-Golgi marker GM130. This result suggests that the cargo did not reach the *cis*-cisternae at 0 min (Figure 3A). However, at 5 min after illumination, VSVG was colocalized with GM130; at 10 min after illumination, some of the cargoes began to reach the *trans*-Golgi cisternae marked by GalT::iRFP713. Furthermore, at 30 min after illumination, the cargoes had completely passed through the *trans-*Golgi cisternae and gradually disappeared. Finally, at 120 min after illumination, no cargoes were detected around the Golgi/RE units. Unexpectedly, when dynasore was administered before cargo release by illumination, the transport of VSVG was stalled at the *cis*-side of the Golgi stacks (Figure 3C). VSVG continuously colocalized strongly with GM130 for at least 45 min (Figure 3D). However, at 120 min after illumination, the cargoes were detected between GM130 and GalT (Figure 3E), indicating that the cargoes were slowly sent forward. Meanwhile, even at 60 min after administration of dynasore, the Golgi stacks strongly maintained their *cis-trans* polarity (Figure 3B). We also observed the effects of dynasore on the transport of GPI-AP and TNFα. After administration of dynasore, both GPI-AP and TNFα continued to colocalize with GM130 even at 90 min and 60 min after illumination respectively, similar to the results observed for VSVG (Figure 3F, G).

**Figure 3.**
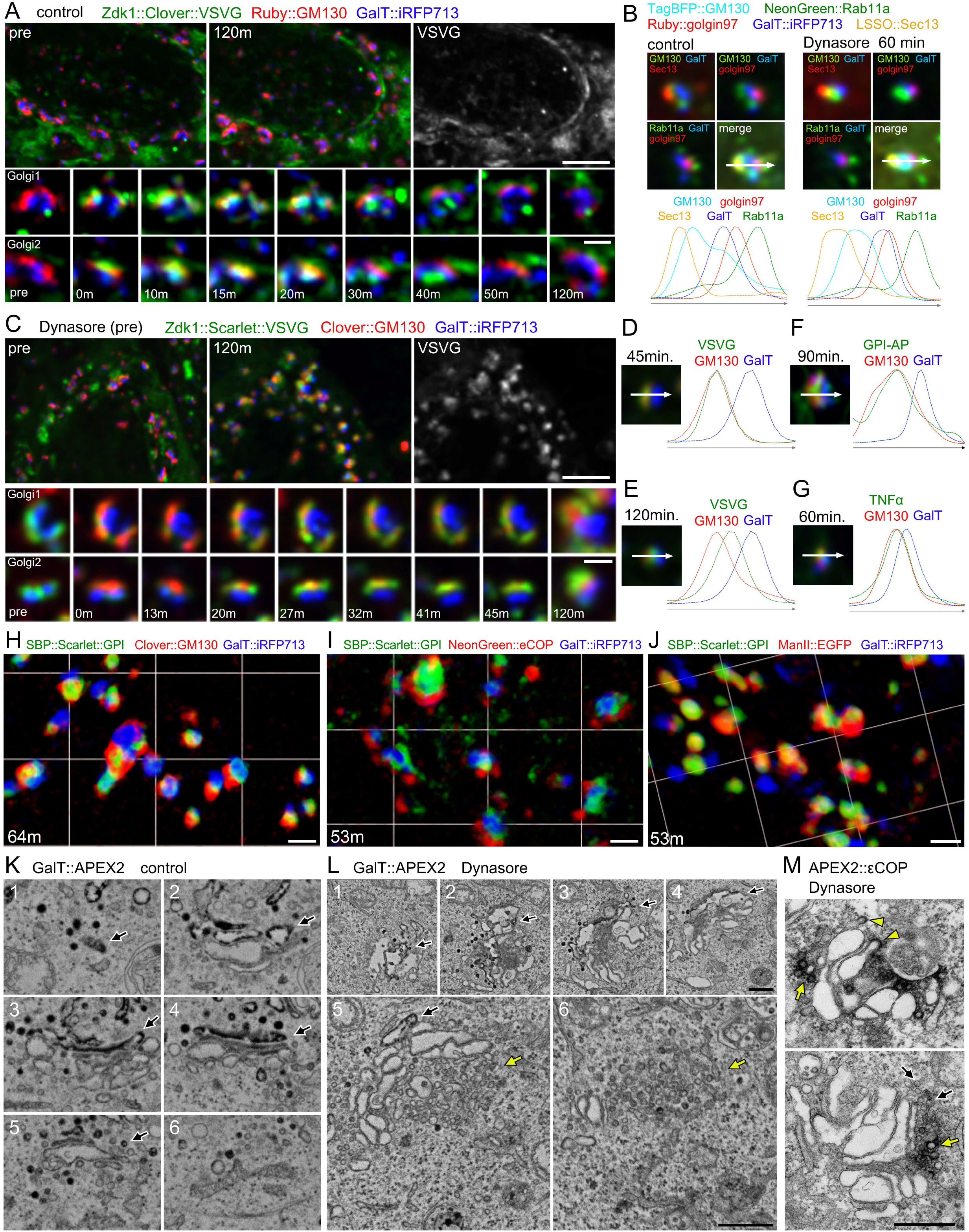
Pre-administration of dynasore-inhibited cargo transport at the *cis*-side of the Golgi stacks. (A) Localization of Zdk1::Clover::VSVG before illumination (upper left) and 120 min after illumination (upper middle and right) in untreated cells. The time time-course course of localization of Zdk1::Clover::VSVG before and after illumination of an untreated single Golgi stack (bottom). Zdk1::Clover::VSVG is shown in green, a *cis*-Golgi marker Ruby::GM130 in red, and a *trans*-Golgi marker GalT::iRFP713 in blue. (B) Upper panels show examples of quintuple-stained Golgi/RE units using the indicated fluorescent tagged proteins before and 1 h after administration of dynasore. Plots show signal intensities from the image on the upper panel. Signal intensity was measured along the arrow (representing 1.5 μm) in the inset. The graph shows the overlap between channels. (C) Localization of Zdk1::Scarlet::VSVG before illumination (upper left) and at 120 min after illumination (upper middle and right) in untreated cells. The time-course of localization of Zdk1::Scarlet::VSVG before and after illumination of the dynasore-treated single Golgi stack (bottom). Zdk1::Scarlet::VSVG is shown in green, *cis*-Golgi marker Clover::GM130 in red, and *trans*-Golgi marker GalT::iRFP713 in blue. (D–G) Plots show signal intensities from the image on the left Golgi/RE unit at 45 min (D), 120 min (E), 90 min (F), and at 60 min (G) after illumination. Signal intensity was measured along the arrow (representing 1.5 μm). The cargoes Zdk1::Scarlet::VSVG (D, E), Zdk1::Clover::GPI (F), and TNFα::Zdk1::Clover (G) are in green, *cis*-Golgi marker Clover/Ruby::GM130 in red, and *trans*-Golgi marker GalT::iRFP713 in blue. (H–J) Volumetrically-rendered images for localization of SBP::Scarlet::GPI (green) in dynasore-administered cells after biotin-addition observed using SCLIM. Clover::GM130 (H), NeonGreen::eCOP (I), or ManII::EGFP (J) is shown in red and *trans*-Golgi marker GalT::iRFP713 in blue. The time shown in the bottom-left corner is the time after biotin-addition. (K, L) Scanning electron micrographs of serial sections of a Golgi stack with 200 nm-interval after 1 h of incubation with (L) or without 100 μM dynasore (K). GalT::APEX2 visualized trans-Golgi cisternae and vesicles (black arrows). Yellow arrows indicate the accumulation of vesicles on the *cis*-side of the Golgi stack. (M) Transmission electron micrographs of Golgi stacks with APEX2::eCOP, which marked COPI budding profiles (yellow arrowheads) and vesicles (yellow arrows). Black arrows indicate vesicles without staining. Scale bars: 5 μm (upper panel in A, C), 1 μm (lower panel in A, C), 2 μm (H–J) and 500 nm (K–M).

We examined the precise localization of cargoes using super-resolution confocal live imaging microscopy (SCLIM). Because of the limitation of the light source equipped on SCLIM, we used the RUSH system for the transport assay rather than using RudLOV. We found that with prior administration of dynasore, the cargoes were partially and fully colocalized with the *cis*-cisternae marker GM130 and *medial*-cisternae marker ManII at 53 and 64 min after biotin addition, respectively. However, these proteins did not colocalize with eCOP (Figure 3H–J). Thus, the cargoes were located within the *cis-* and *medial*-cisternae rather than in the COPI vesicles. These results suggest that dynasore inhibits the maturation of *cis*/*medial-*Golgi cisternae to *trans-*Golgi cisternae.

Ultrastructural observation of dynasore-treated cells using electron microscopy revealed that the Golgi stacks were accompanied by numerous vesicles (Figure 3K, L and Video 1). These vesicles (Figure 3L yellow arrows) were mainly found on the opposite side to the GalT::APEX2-marked *trans-*Golgi cisternae (Figure 3L, black arrows), suggesting that the vesicles localized around the *cis*-side of the Golgi stacks. Moreover, as some vesicles were positively marked by eCOPI::APEX2 (Figure 3M, yellow arrows), these vesicles were likely coated with COPI. These results suggest that dynasore inhibits the uncoating, tethering, or fusion of COPI vesicles to prevent the maturation of *cis*/*medial-*Golgi cisternae to *trans-*Golgi cisternae. Off target effects of dynasore on fluid-phase endocytosis and membrane ruffling has been reported (Park *et al*, 2013), but there is no report showing inhibition of early Golgi transport by dynasore, as we found here.

### Dynasore additionally exerted a canonical inhibitory effect on cargo export from the TGN

The unexpected inhibition of early Golgi cargo transport by dynasore can mask its canonical inhibitory effect on dynamin-2 in the TGN. To prevent this effect, we delayed dynasore administration to slightly after cargo release and evaluated various intervals between the times of illumination and dynasore administration. Notably, administration of dynasore at 4 min after illumination allowed VSVG to pass through the *cis*- and *trans*-Golgi cisternae but inhibited export from the Golgi/RE units. VSVG remained at the supposed TGN on the *trans* side of the GalT::iRFP713 compartment for more than 120 min following illumination (Figure 4A–C). Similar post-illumination administration was performed for GPI-AP and TNFα. We observed similar *cis* to *trans* migration of the cargoes and inhibition of exit from the TGN (Figure 4D, E). The distinct effects of pre- and post-illumination administration of dynasore were also observed in cells without nocodazole treatment (Figure S1E–H).

**Figure 4.**
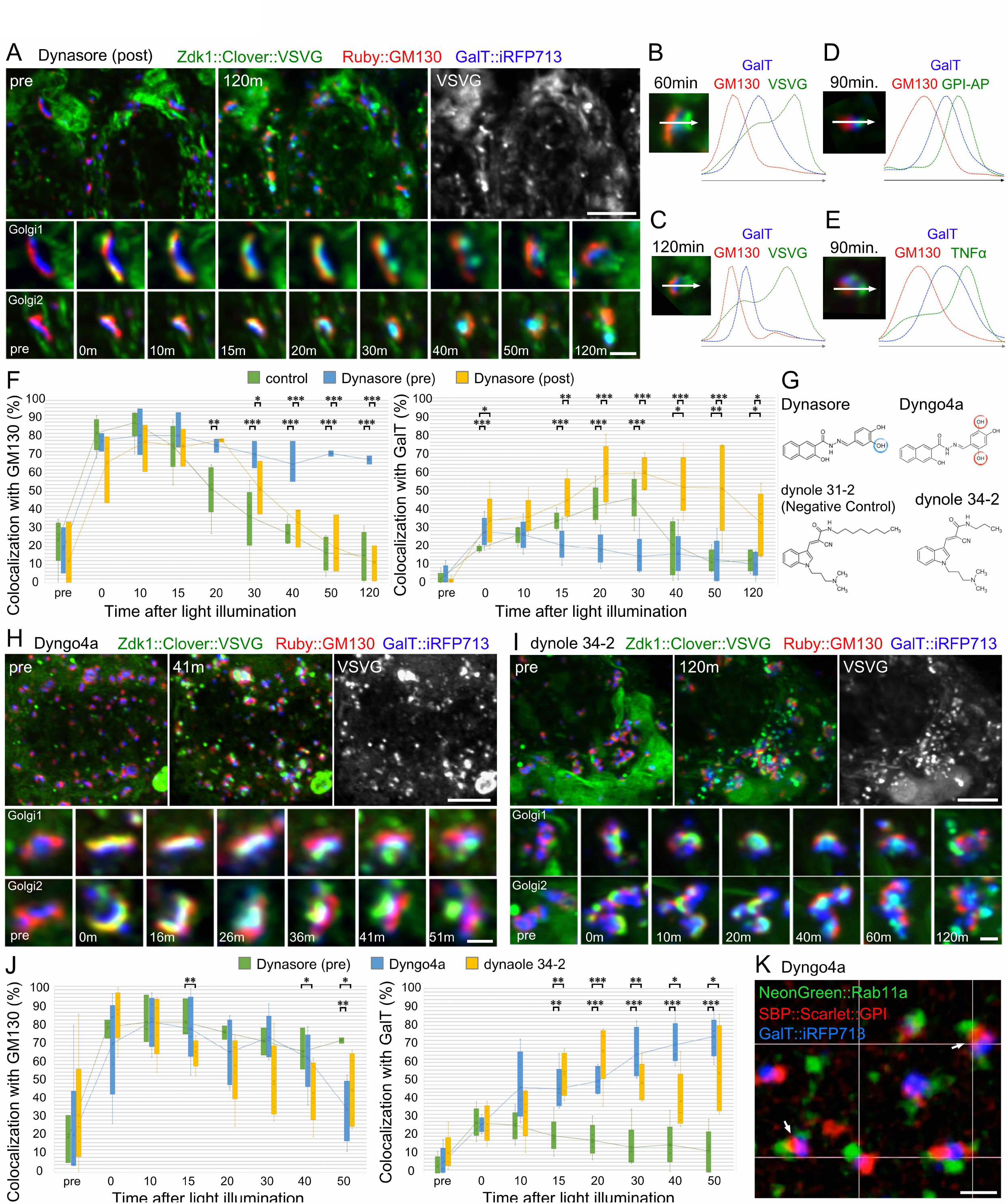
Post-administration of dynasore at 4 min after illumination specifically inhibits cargo transport at the *trans*-Golgi network (TGN) (A) Localization of Zdk1::Clover::VSVG before illumination (upper left) and 120 min after illumination (upper middle and right) in cells post-administered with dynasore at 4 min after illumination. The time-course of localization of Zdk1::Clover::VSVG before and after illumination in a single Golgi stack (bottom) with post-administered dynasore. Zdk1::Clover::VSVG is shown in green, *cis*-Golgi marker Ruby::GM130 in red, and *trans*-Golgi marker GalT::iRFP713 in blue. (B–E) Plots show signal intensities from the image on the left Golgi/RE unit at 60 min (B), 120 min (C), 90 min (D), and 90 min (E) after illumination of cells with post-administration of dynasore at 4 min after illumination. Signal intensity was measured along the region indicated by the arrow (representing 1.5 μm). The cargoes, Zdk1::Clover::VSVG (B, C), Zdk1::Clover::GPI (D), and TNFα::Zdk1::Clover (E) are in green; *cis*-Golgi marker Ruby::GM130 in red; and *trans*-Golgi marker GalT::iRFP713 in blue. (F) Plots show the colocalization of cargoes with Ruby::GM130 (left) and GalT::iRFP713 (right) in cells that are untreated (green), pre-administered (blue), and post-administered (yellow) with dynasore. Error bars indicate the standard deviation. ***P < 0.001; **P < 0.01; *P < 0.05 (two-tailed unpaired Student’s *t*-test). (G) Structure formulas of four compounds used in this work. Dyngo4a has an extra hydroxy group compared with Dynasore. (H, I) Localization of Zdk1::Clover::VSVG before illumination (upper left) and at 120 min after illumination (upper middle and right) in cells pre-administered with Dyngo4a (H) or dynole 34-2 (I). The time-course of localization of Zdk1::Clover::VSVG before and after illumination in Dyngo4a-treated or (H) or dynole 34-2-treated (I) single Golgi stack (bottom). Zdk1::Clover::VSVG is in green, *cis*-Golgi marker Ruby::GM130 in red, and *trans*-Golgi marker GalT::iRFP713 in blue. (J) Plots showing colocalization of cargoes with Ruby::GM130 (left) and GalT::iRFP713 (right) in cells pre-administered with dynasore (green), Dyngo4a (blue), or dynole 34-2 (yellow). Error bars indicate the standard deviation. ***P < 0.001; **P < 0.01; *P < 0.05 (two-tailed unpaired Student’s *t*-test). (K) Volumetrically-rendered images for localization of SBP::Scarlet::GPI (red) in Dyngo4a-administered cells 35 min after biotin-addition observed using SCLIM. The RE marker NeonGreen::Rab11a is in green and *trans*-Golgi marker GalT::iRFP713 is in blue. Arrows indicate the localization of SBP::Scarlet::GPI on the membrane between trans-Golgi cisternae and RE. Scale bars: 5 μm (upper panel in A, H, I), 1 μm (lower panel in A, H, I), and 2 μm (K).

The time points of co-localization of VSVG with GM130 and GalT in dynasore-administered cells before and after illumination and in dynasore-untreated control cells were plotted (Figure 3F). GM130/VSVG-colocalization peaked at approximately 10 min after illumination in untreated cells. GalT/VSVG co-localization gradually increased, peaking at approximately 30 min after illumination, and then decreased quickly. In pre-illumination dynasore-administered cells, GM130/VSVG-colocalization remained high at 120 min after illumination, and GalT/VSVG-colocalization was not elevated during the 120 min observation period. In post-illumination dynasore-administered cells, after temporal GM130/VSVG colocalization, GalT/VSVG co-localization reached a peak at 20 min and remained high until at least 50 min. These plots clearly indicate that intra-Golgi transport was inhibited by dynasore at the early stage but not at the later stage by pre-illumination administration of dynasore.

### Other dymanin inhibitors, Dyngo4a and dynole 34-2, inhibit cargo transport only at the TGN

The inhibition of exit from the TGN agrees with the results of previous studies, which showed that dynamin-2 is required for forming post-Golgi vesicles in the TGN (Cao *et al*., 2005; Jones *et al*., 1998; Kessels *et al*., 2006; Kreitzer *et al*., 2000; Salvarezza *et al*., 2009). However, inhibition of the early stages of transport in the Golgi stacks by dynasore was unexpected. Thus, we examined the effects of other dynamin inhibitors, such as Dyngo4a, dynole 34-2, and dynole31-2 (negative control) (McCluskey *et al*, 2013; Robertson *et al*, 2014; Stallaert *et al*, 2018). Pre-illumination administration of either Dyngo4a or dynole 34-2 inhibited cargo export from the TGN but did not strongly affect early Golgi transport. Pre-administration of dynole31-2 did not affect cargo transport at all. The time-course plots of GM130/VSVG- or GalT/VSVG-colocalization showed that pre-illumination administration of Dyngo4a or dynole 34-2 allowed cargo migration from GM130-positive cisternae to GalT-positive cisternae, whereas exit from the Golgi/RE units was inhibited. As GPI-AP is transported to the GA-RE after passing through GalT-positive *trans*-cisternae in untreated cells, we investigated whether GPI-AP reached the GA-RE in Dyngo4a-treated cells. Observations using SCLIM clearly revealed that GPI-AP was separated from the RE marker Rab11a. Thus, Dyngo4a inhibits transport from the Golgi stacks to the GA-RE.

These effects of Dyngo4a and dynole 34-2 agree with the reports that dynamin-2 is essential for post-Golgi transport but not for early Golgi transport (Cao *et al*., 2005; Jones *et al*., 1998; Kessels *et al*., 2006; Kreitzer *et al*., 2000; Salvarezza *et al*., 2009). Thus, dynasore likely inhibits dynamin-2 at the TGN, resulting in inhibition of post-Golgi transport; it additionally inhibits another target at the *cis*-side of Golgi stacks, resulting in defects in the maturation of *cis*/*medial*-cisternae to *trans-*cisternae. Cisternal maturation of the Golgi stack is driven by retrograde trafficking of Golgi-resident proteins by COPI vesicles (Emr *et al*, 2009; Ishii *et al*, 2016; Papanikou *et al*, 2015; Papanikou & Glick, 2014). Together with the electron micrographs showing COPI vesicle accumulation in dynasore-administered cells, the unidentified target of dynasore at the *cis*-side of the Golgi stacks may be involved in uncoating/tethering/fusion of COPI vesicles.

Our results indicate that cargo transport in the early (*cis*/*medial* to *trans*), but not late (*trans* to TGN), stages is inhibited by dynasore. Thus, dynasore may inhibit the factors involved in early cisternal maturation. Interestingly, the tether and SNAREs involved in the fusion of COPI vesicles with the ER differ from those involved in the fusion of COPI vesicles with Golgi cisternae; the Dsl1 complex and Sec20/Ufe1/Use1/Sec22 are associated with the former, whereas the COG complex and Stx5/Gos28/Ykt6/GS15 are associated with the latter (Hong & Lev, 2014; Ren *et al*, 2009; Zink *et al*, 2009). Additionally, there are three types of ARFs with different localizations on the *cis*-side of Golgi stacks: ARF4/5 is mainly localized on the ERGIC, and ARF1 is mainly localized on the *cis*-Golgi cisternae and TGN (Chun *et al*, 2008; Duijsings *et al*, 2009; Wong-Dilworth *et al*, 2023) (Chun *et al*., 2008; Duijsings *et al*., 2009; Wong-Dilworth *et al*., 2023). The function of ARF4/5 is not fully understood but they have the activity for COPI assembly at least in vitro (Adolf *et al*, 2019; Popoff *et al*, 2011). ARF1 is well-known to be required for not only assembly, but also scission and uncoating of COPI vesicles; scission of COPI vesicles from Golgi-cisternae requires ARF1 dimers (Beck *et al*, 2011; Diestelkoetter-Bachert *et al*, 2020), and GTP-hydrolysis on ARF1 is pre-requisite for COPI uncoating (Tanigawa *et al*, 1993; Zink *et al*., 2009). Recent studies indicated that retrograde trafficking to the ER is impaired in ARF4 knockout but not in ARF1 knockout cells (Pennauer *et al*, 2022). Yeast cells house two ARFs, namely, Arf1 and Arf2, and it is difficult to tell which yeast Arf is the functional counterpart of mammalian ARF4/5 based on homology analysis. Notably, the maturation of early Golgi cisternae becomes slower and less frequent in Arf1-deficient yeasts, whereas the maturation of late Golgi cisternae does not change (Bhave *et al*, 2014). Thus, dynasore may inhibit GTP-hydrolysis on ARF4/5 or COPI-uncoating/tethering/fusion machinery in the early-Golgi cisternae or ER but not in the late-Golgi cisternae. Future studies are required to identify dynasore targets on the *cis*-side of the Golgi stacks to reveal the underlying mechanism of cisternal maturation.

## Conclusion

We developed a new cargo releasing method, RudLOV, which has three exceptional control abilities: spatial, temporal, and quantitative control of cargo release. RudLOV’s quantitative control of cargo release enables visualization of cisternal movement of cargos and the cargo-specific exit site on the Golgi/TGN. Moreover, RudLOV’s precise temporal control ability of cargo release can be used to dissect canonical and non-canonical effects of the well-known dynamin inhibitor dynasore. LOV2 can be activated by 445 or 488nm light offering damage-less application of light-induced cargo export. Moreover, the simplicity of RudLOV without chemical application allows for cargo export in cells *in situ* or in whole animals. These exceptional control abilities and convenience of use demonstrate the potential of using RudLOV for sophisticated observation of cargo trafficking.

## Materials and Methods

### Construction of plasmids for RudLOV and organelle markers

The fluorescent tags described in this paper, Clover, Ruby, NeonGreen and Scarlet, are tandem repeat of two identical fluorescent proteins, mClover3, mRuby3 (gift from Michael Lin (Bajar *et al*, 2016)), mNeonGreen and mScarlet-I (gift from Erik Dent (Taylor *et al*, 2020)), respectively. Details of plasmid vectors used in this paper are described in supplemental table 1. Sequence files in GenBank format are available in Dryad (https://doi.org/10.5061/dryad.jm63xsjhs).

### Stably transformed HeLa cells expressing LOV2 hook—hSec61b::tagBFP2::LOV2

HeLa cells stably expressing GalT::iRFP713 (clone C9) (Fujii *et al*., 2020a) were transfected with pEBP-Sec61b-mTagBFP2-LOV2 using jetPRIME transfection reagent (Polyplus-transfection, Illkirch-Graffenstaden, France). After 24 h, the cells were selected using 2 µg/mL of puromycin (Fujifilm WAKO Chemicals, Osaka, Japan). The clones of stable transformants were established using limited dilutions. A clone showing strong tagBFP2 expression was selected for further experiments (clone G7).

### Transferrin uptake in HeLa cells

After 4 h of treatment with 10 μM nocodazole (Cayman Chemical, Ann Arber, USA), GalT::iRFP713 expressing HeLa cells (Clone C9) were preincubated for 30 min in serum-free medium containing nocodazole. The cells were then incubated with 30 μg/mL of Alexa Fluor 568 conjugated Tfn (Tfn-568) (Life Technologies, Carlsbad, CA, USA) for 5 min and chased for 8 min in nocodazole-containing medium. Dynasore (100 μM) (Cayman Chemical) was added to the serum-free medium, medium with Tfn, and chasing medium.

### Live imaging of HeLa cells using RudLOV by FV3000

HeLa cells stably expressing GalT::iRFP713 and Sec61b::TagBFP2::LOV2 (clone G7) were transfected with a DNA plasmid encoding Zdk1-FP-GPI, Zdk1-FP-VSVG or TNFα-Zdk1-FP using jetPRIME or JetOptimus transfection reagent according to the manufacturer’s instructions. At 6–10 h after transfection, the medium was replaced with fresh medium containing 25 µM of biliverdin (Cayman Chemical). The next day, the cells were treated for 4 h with nocodazole. Cycloheximide (100 μg/mL) (Cayman Chemical) was administered, and Zdk1-FP-cargo release into the secretory pathway was induced by administration of 0.05% 445 nm laser illumination for 5–10 min using the LSM stimulation mode in FV3000 (Evident, Tokyo, Japan), equipped with a quinta Band dichroic mirror 89903bs (Chroma Technology Corporation, Bellows Falls, VT, USA). Time lapse 3D-stacks of confocal micrographs were obtained using an FV3000 (except for those shown in Figure 1F). The 3D-stacks were projected using the maximum intensity with Fiji. The obtained 3D voxel data were displayed as volume-rendered images using Volocity software. For Figure 1F, a single section of the confocal micrograph for pre-illumination or 15 min after-illumination condition was obtained using an FV3000. As shown in Figure 1G, confocal micrographs were obtained, and the signal intensity of Zdk1-FP-cargoes was measured in the Golgi area defined by GalT::iRFP713 using Fiji. Precise cargo localization within the Golgi stacks was measured using line profiles across the Golgi stacks using Fiji and processed using Plot2 Pro (Michael Wesemann, Berlin, Germany). Co-localization of VSVG and GM130/GalT was measured using custom plugins for Fiji, as described by Papanikou et al. (Papanikou *et al*., 2015).

We administered 100 μM of dynasore, 10 μM of Dyngo4a (Cayman Chemical), 10 μM of dynole34-2, and 100 μM of dynole31-2 (negative control) (R&D Systems, Minneapolis, MN, USA) before or after illumination, as indicated in the text.

### Live imaging of HeLa cells using the RUSH system with an FV3000 confocal microscope

HeLa cells stably expressing GalT::iRFP713 were transfected with a DNA plasmid encoding RUSH system bi-cistronic expression plasmids Str::KDEL_SBP::NeonGreen::VSVG using jetPRIME or JetOptimus transfection reagent as per the manufacturer’s instructions. At 6–10 h after transfection, the medium was replaced with fresh medium containing 25 µM of biliverdin. The next day, after 4 h of treatment with nocodazole, the release of SBP::NeonGreen::VSVG into the secretory pathway was induced by replacing the medium with 100 µM biotin (Fujifilm WAKO Chemicals) and 100 μg/mL of cycloheximide. Time-lapse confocal micrographs were obtained using an FV3000 microscope. As shown in Figure 1E, the signal intensity of SBP::Neon Green::VSVG was measured in proximity to the Golgi using Fiji as defined by GalT::iRFP713.

### Live imaging of HeLa cells using the RUSH system with SCLIM

HeLa cells stably expressing GalT::iRFP713 inoculated on glass-based dishes (Iwaki, Tolyo, Japan) were transfected with a DNA plasmid encoding the RUSH system bi-cistronic expression plasmids Str::KDEL_SBP::EGFP::GPI using JetOptimus transfection reagent as per the manufacturer’s instructions. The cells were cultured for one day in phenol red-free medium to reduce background fluorescence. The cells were then treated with 10 µM nocodazole for 4 h to disrupt the microtubules. SBP::EGFP::GPI was released into the secretory pathway by replacing the medium with 100 µM biotin and 100 μg/mL of cycloheximide. To observe the effect of Dyngo4a, the cells were pre-treated with 100 µM of Dyngo4a for approximately 10 min. The medium was replaced with medium containing 100 µM biotin, 100 μg/mL cycloheximide, and 100 µM Dyngo4a to initiate the transport of SBP::EGFP::GPI. Z-stack images were obtained using SCLIM (Kurokawa *et al*, 2013; Kurokawa *et al*., 2019; Tojima *et al*, 2023) and processed via deconvolution with Volocity (Perkin Elmer, Waltham, MA, USA) using the theoretical point-spread function for spinning-disk confocal microscopy. The obtained 3D voxel data were displayed as volume-rendered images using Volocity software.

### Electron microscopy imaging of GalT::APEX2::EGFP expressing HeLa cells with and without dynasore treatment

Aclar-film (Nisshin-EM, Tokyo, Japan) was cut into pieces, washed with acetone, placed in an µ-Slide 8-well cell culture chamber (ibidi GmbH, Gräfelfing, Germany), and hydrated using culture medium for more than 24 h. HeLa cells that were transfected with CMV-GalT::APEX2::EGFP or CT7-NeonGreen::APEX2::eCOP were seeded on the film and grown overnight. After 1 h of incubation with 100 μM of dynasore (Cayman Chemical), the cells were fixed in 0.1 M cacodylate buffer (pH 7.4) with 2% glutaraldehyde (Electron Microscopy Sciences, Hatfield, PA, USA) and 2% paraformaldehyde (Electron Microscopy Sciences) for 1 h on ice. DAB staining was performed as described by Martell et al. (Martell *et al*, 2017) with some previously described modifications (Otsuka *et al*, 2019). DAB staining was performed by applying fresh DAB solution every 20 min for 1 h at room temperature to cells expressing GalT::APEX2::EGFP and eCOP::APEX2::EGFP. Post-fixation was performed using 2% (w/v) osmium tetroxide (Electron Microscopy Sciences, Hatfield, UAS) for 30 min in chilled cacodylate buffer. The films were rinsed five times (2 min each time) with chilled distilled water, placed overnight in ddH_2_O containing chilled 2% (w/v) uranyl acetate, dehydrated, and penetrated with EPON-812 resin (Electron Microscopy Sciences) as described previously (Otsuka *et al*., 2019). Aclar-films with cells in EPON-812 were polymerized into a thin EPON-812 sheet at 100 °C for 20 h. Polymerized EPON-812 sheet-containing cells were cut into 2 mm squares, and the attached Aclar-film was removed and re-polymerized after being placed on the bottom of a pyramidal mold with new EPON-812. The cells were cut with a diamond knife into 70-nm sections and imaged using a JEM1400 transmission electron microscope (JEOL, Tokyo, Japan) operated at 80 kV. Montage images were captured with a CCD camera system (JEOL).

### Serial section scanning electron microscopy observation of GalT::APEX2::EGFP-expressing HeLa cells

Serial section scanning electron microscopy was performed using a high-resolution field-emission scanning electron microscope and back-scattered electron detector. Serial ultrathin sections (thickness of 50 nm) were cut using a diamond knife (Diatome 45°) on an ultramicrotome (EM UC7, Leica Microsystems, Wetzlar, Germany) and placed on silicon wafers (10 × 22 mm). The sections were stained with 0.4% uranyl acetate for 10 min and lead stain solution (Sigma-Aldrich, St. Louis, MO, USA) for 2 min and coated with osmium tetroxide using an osmium coater (HPC-1SW, Vacuum Device Inc., Mito, Japan). Serial sections were observed using a field-emission scanning electron microscope (Regulus8240; Hitachi High-Tech, Tokyo, Japan) equipped with auto-capture for array tomography and low-angle back-scattered electron detector at an accelerating voltage of 2 kV.

## Author Contributions

T.S. and A.S. designed the study; T.O. performed the initial experiments to develop RudLOV. T.T.^1^ performed most of the laboratory experiments and analyzed the data; Y.G. and K.T. performed serial sectioning and observation of the cells expressing GalT::APEX2. T.T.^3^ and A.N. supervised SCLIM observation and data analyses. T.S. supervised the molecular biology experiments. A.S. supervised all aspects of the project. T.S. and A.S. wrote the paper with input and final approval from T.T^1^., T.O., Y.G., K.T., T.T.^3^, and A.N.

## Acknowledgements

This work was supported by the Japan Society for the Promotion of Science (JSPS) (KAKENHI grant no. 22H02617) to A.K.S., (KAKENHI grant no. 19K06566) to T.S., (KAKENHI grant no. 22K06213) to T.T^3^ and (KAKENHI grants no. 18H05275 and 23H00382) to A.N.; Japan Science and Technology Agency (JST) (PRESTO grant no. 25-J-J4215 and CREST grant no. JPMJCR22E2) to A.K.S, (CREST grant no. JPMJCR21E3) to T.T^3^ and (SPRING grant no. JPMJSP2132) to T.T.^1^; and Core Research for Organelle Diseases funding from Hiroshima University, Takeda Science Foundation and Ohsumi Frontier Science Foundation to A.K.S. Grant Number JP22H04926, Grant-in-Aid for Transformative Research Areas ― Platforms for Advanced Technologies and Research Resources “Advanced Bioimaging Support” to K.T. We thank Rumi Sato and Takaharu Okada (RIKEN IMS) for offering the FE-SEM equipped with auto-capture for array tomography system. We would like to thank Editage (http://www.editage.com) for editing and reviewing this manuscript for English language"

## Figure legends

**Figure S1. Dynasore inhibits cargo transport at the *cis*- or *trans*-side of Golgi stacks in nocodazole-untreated cells**

Cargoes are shown in green and GalT::iRFP713 in magenta.

(A, B) Localization of Zdk1::Clover::GPI before (left) and at 0, 5, 10, 20, 40, and 60 min after illumination at 445 nm using the RudLOV system in untreated (A) and nocodazole-treated cells (B). Inset in B shows a magnified image of Zdk1::Clover::GPI. Cargoes are shown in green and GalT::iRFP713 in magenta.

(C, D) Localizations of TNFα::Zdk1::Clover before (left) and at 0, 5, 10, 20, 40, 50 min after illumination with 445 nm using the RudLOV system in untreated (C) and nocodazole-treated cells (D). Inset in D shows the magnified image of TNFα::Zdk1::Clover. Cargoes are shown in green and GalT::iRFP713 in blue.

(E) Localization of Zdk1::Scarlet::VSVG before illumination and at 0, 37, and 118 min after illumination in cells pre-treated with dynasore. The cargoes are in green, *cis*-Golgi marker Clover::GM130 in red, and *trans*-Golgi marker GalT::iRFP713 in blue.

(F) Plots showing signal intensities from the image of the left Golgi/RE unit ast 98 min after illumination in the cell pre-administered with dynasore. Signal intensity was measured along the arrow (representing 3 μm).

(G) Localization of Zdk1::Scarlet::VSVG before illumination and at 0, 43, and 124 min after illumination in cells that were post-administrated dynasore 4 min after illumination. The cargoes are in green, *cis*-Golgi marker Clover::GM130 in red, and *trans*-Golgi marker GalT::iRFP713 in blue.

(H) Plots show signal intensities from the image of the left Golgi/RE unit at 97 min after illumination in dynasore pre-administered cell. Signal intensity was measured along the arrow (representing 3 μm).

(I) Uptake of Tfn in untreated and dynasore-treated cells at 8 min after incubation with 30 μg/ml of Alexa Fluor 568-conjugated Tfn.

(J) Localization of Zdk1::Clover::VSVG before illumination (upper left) and 60 min after illumination (upper middle and right) in cells pre-treated with dynole 31-2 (negative control: NC). Time-course of Zdk1::Clover::VSVG localization before and after illumination in a dynole NC-treated single Golgi stack (bottom). Zdk1::Clover::VSVG is in green, *cis*-Golgi marker Ruby::GM130 in red, and *trans*-Golgi marker GalT::iRFP713 in blue.

Scale bars: 5 μm (A–E), 2 μm (insets in B, D), 5 μm (G), 10μm (I), 5 μm (upper panel J), and 1μm (downer panel J).

---

**Video 1. Accumulation of vesicles at the *cis*-side of Golgi/RE unit in dynasore-treated HeLa cells**

Video 1 contains two movies built from scanning electron micrographs for 50 nm-thick serial sections of Golgi/RE units in HeLa cells untreated and treated 1 h with dynasore. *Trans*-Golgi cisterna was visualized with GalT::APEX2 as electron-dense material.

**Table S1. Information of markers and plasmids used in this work**

Backbone and inserts with coding region are described. For further information, electronic files in GenBank format in available in Dryad (https://doi.org/10.5061/dryad.jm63xsjhs)

## References

Adolf F, Rhiel M, Hessling B, Gao Q, Hellwig A, Béthune J, Wieland FT (2019) Proteomic Profiling of Mammalian COPII and COPI Vesicles. Cell Rep 26: 250–265.e255

Bajar BT, Wang ES, Lam AJ, Kim BB, Jacobs CL, Howe ES, Davidson MW, Lin MZ, Chu J (2016) Improving brightness and photostability of green and red fluorescent proteins for live cell imaging and FRET reporting. Sci Rep 6: 20889

Beck R, Prinz S, Diestelkötter-Bachert P, Röhling S, Adolf F, Hoehner K, Welsch S, Ronchi P, Brügger B, Briggs JA et al (2011) Coatomer and dimeric ADP ribosylation factor 1 promote distinct steps in membrane scission. J Cell Biol 194: 765–777

Bhave M, Papanikou E, Iyer P, Pandya K, Jain BK, Ganguly A, Sharma C, Pawar K, Austin J, Day KJ et al (2014) Golgi enlargement in Arf-depleted yeast cells is due to altered dynamics of cisternal maturation. J Cell Sci 127: 250–257

Boncompain G, Divoux S, Gareil N, de Forges H, Lescure A, Latreche L, Mercanti V, Jollivet F, Raposo G, Perez F (2012) Synchronization of secretory protein traffic in populations of cells. Nat Methods 9: 493–498

Bourke AM, Schwartz SL, Bowen AB, Kleinjan MS, Winborn CS, Kareemo DJ, Gutnick A, Schwarz TL, Kennedy MJ (2021) zapERtrap: A light-regulated ER release system reveals unexpected neuronal trafficking pathways. J Cell Biol 220

Cao H, Weller S, Orth JD, Chen J, Huang B, Chen JL, Stamnes M, McNiven MA (2005) Actin and Arf1-dependent recruitment of a cortactin-dynamin complex to the Golgi regulates post-Golgi transport. Nat Cell Biol 7: 483–492

Casler JC, Zajac AL, Valbuena FM, Sparvoli D, Jeyifous O, Turkewitz AP, Horne-Badovinac S, Green WN, Glick BS (2020) ESCargo: a regulatable fluorescent secretory cargo for diverse model organisms. Mol Biol Cell 31: 2892–2903

Chun J, Shapovalova Z, Dejgaard SY, Presley JF, Melançon P (2008) Characterization of class I and II ADP-ribosylation factors (Arfs) in live cells: GDP-bound class II Arfs associate with the ER-Golgi intermediate compartment independently of GBF1. Mol Biol Cell 19: 3488–3500

Diestelkoetter-Bachert P, Beck R, Reckmann I, Hellwig A, Garcia-Saez A, Zelman-Hopf M, Hanke A, Nunes Alves A, Wade RC, Mayer MP et al (2020) Structural characterization of an Arf dimer interface: molecular mechanism of Arf-dependent membrane scission. FEBS Lett 594: 2240–2253

Duijsings D, Lanke KH, van Dooren SH, van Dommelen MM, Wetzels R, de Mattia F, Wessels E, van Kuppeveld FJ (2009) Differential membrane association properties and regulation of class I and class II Arfs. Traffic 10: 316–323

Emr S, Glick BS, Linstedt AD, Lippincott-Schwartz J, Luini A, Malhotra V, Marsh BJ, Nakano A, Pfeffer SR, Rabouille C et al (2009) Journeys through the Golgi--taking stock in a new era. J Cell Biol 187: 449–453

Farr GA, Hull M, Mellman I, Caplan MJ (2009) Membrane proteins follow multiple pathways to the basolateral cell surface in polarized epithelial cells. J Cell Biol 186: 269–282

Ferguson SM, De Camilli P (2012) Dynamin, a membrane-remodelling GTPase. Nat Rev Mol Cell Biol 13: 75–88

Ferguson SM, Raimondi A, Paradise S, Shen H, Mesaki K, Ferguson A, Destaing O, Ko G, Takasaki J, Cremona O et al (2009) Coordinated actions of actin and BAR proteins upstream of dynamin at endocytic clathrin-coated pits. Dev Cell 17: 811–822

Fourriere L, Divoux S, Roceri M, Perez F, Boncompain G (2016) Microtubule-independent secretion requires functional maturation of Golgi elements. J Cell Sci 129: 3238–3250

Fujii S, Kurokawa K, Inaba R, Hiramatsu N, Tago T, Nakamura Y, Nakano A, Satoh T, Satoh AK (2020a) Recycling endosomes attach to the trans-side of Golgi stacks in. J Cell Sci

Fujii S, Tago T, Sakamoto N, Yamamoto T, Satoh T, Satoh AK (2020b) Recycling endosomes associate with Golgi stacks in sea urchin embryos. Commun Integr Biol 13: 59–62

Hong W, Lev S (2014) Tethering the assembly of SNARE complexes. Trends Cell Biol 24: 35–43

Ishii M, Suda Y, Kurokawa K, Nakano A (2016) COPI is essential for Golgi cisternal maturation and dynamics. J Cell Sci 129: 3251–3261

Jones SM, Howell KE, Henley JR, Cao H, McNiven MA (1998) Role of dynamin in the formation of transport vesicles from the trans-Golgi network. Science 279: 573–577

Kessels MM, Dong J, Leibig W, Westermann P, Qualmann B (2006) Complexes of syndapin II with dynamin II promote vesicle formation at the trans-Golgi network. J Cell Sci 119: 1504–1516

Kreitzer G, Marmorstein A, Okamoto P, Vallee R, Rodriguez-Boulan E (2000) Kinesin and dynamin are required for post-Golgi transport of a plasma-membrane protein. Nat Cell Biol 2: 125–127

Kurokawa K, Ishii M, Suda Y, Ichihara A, Nakano A (2013) Live cell visualization of Golgi membrane dynamics by super-resolution confocal live imaging microscopy. Methods Cell Biol 118: 235–242

Kurokawa K, Okamoto M, Nakano A (2014) Contact of cis-Golgi with ER exit sites executes cargo capture and delivery from the ER. Nat Commun 5: 3653

Kurokawa K, Osakada H, Kojidani T, Waga M, Suda Y, Asakawa H, Haraguchi T, Nakano A (2019) Visualization of secretory cargo transport within the Golgi apparatus. J Cell Biol 218: 1602–1618

Lippincott-Schwartz J, Roberts TH, Hirschberg K (2000) Secretory protein trafficking and organelle dynamics in living cells. Annu Rev Cell Dev Biol 16: 557–589

Macia E, Ehrlich M, Massol R, Boucrot E, Brunner C, Kirchhausen T (2006) Dynasore, a cell-permeable inhibitor of dynamin. Dev Cell 10: 839–850

Martell JD, Deerinck TJ, Lam SS, Ellisman MH, Ting AY (2017) Electron microscopy using the genetically encoded APEX2 tag in cultured mammalian cells. Nat Protoc 12: 1792–1816

McCluskey A, Daniel JA, Hadzic G, Chau N, Clayton EL, Mariana A, Whiting A, Gorgani NN, Lloyd J, Quan A et al (2013) Building a better dynasore: the dyngo compounds potently inhibit dynamin and endocytosis. Traffic 14: 1272–1289

Otsuka Y, Satoh T, Nakayama N, Inaba R, Yamashita H, Satoh AK (2019) Parcas is the predominant Rab11-GEF for rhodopsin transport in. J Cell Sci 132

Papanikou E, Day KJ, Austin J, Glick BS (2015) COPI selectively drives maturation of the early Golgi. Elife 4

Papanikou E, Glick BS (2014) Golgi compartmentation and identity. Curr Opin Cell Biol 29: 74–81

Park RJ, Shen H, Liu L, Liu X, Ferguson SM, De Camilli P (2013) Dynamin triple knockout cells reveal off target effects of commonly used dynamin inhibitors. J Cell Sci 126: 5305–5312

Pennauer M, Buczak K, Prescianotto-Baschong C, Spiess M (2022) Shared and specific functions of Arfs 1-5 at the Golgi revealed by systematic knockouts. J Cell Biol 221

Popoff V, Langer JD, Reckmann I, Hellwig A, Kahn RA, Brügger B, Wieland FT (2011) Several ADP-ribosylation factor (Arf) isoforms support COPI vesicle formation. J Biol Chem 286: 35634–35642

Presley JF, Cole NB, Schroer TA, Hirschberg K, Zaal KJ, Lippincott-Schwartz J (1997) ER-to-Golgi transport visualized in living cells. Nature 389: 81–85

Ren Y, Yip CK, Tripathi A, Huie D, Jeffrey PD, Walz T, Hughson FM (2009) A structure-based mechanism for vesicle capture by the multisubunit tethering complex Dsl1. Cell 139: 1119–1129

Robertson MJ, Deane FM, Robinson PJ, McCluskey A (2014) Synthesis of Dynole 34-2, Dynole 2–24 and Dyngo 4a for investigating dynamin GTPase. Nat Protoc 9: 851-870

Salvarezza SB, Deborde S, Schreiner R, Campagne F, Kessels MM, Qualmann B, Caceres A, Kreitzer G, Rodriguez-Boulan E (2009) LIM kinase 1 and cofilin regulate actin filament population required for dynamin-dependent apical carrier fission from the trans-Golgi network. Mol Biol Cell 20: 438–451

Shomron O, Nevo-Yassaf I, Aviad T, Yaffe Y, Zahavi EE, Dukhovny A, Perlson E, Brodsky I, Yeheskel A, Pasmanik-Chor M et al (2021) COPII collar defines the boundary between ER and ER exit site and does not coat cargo containers. J Cell Biol 220

Solinger JA, Rashid HO, Prescianotto-Baschong C, Spang A (2020) FERARI is required for Rab11-dependent endocytic recycling. Nat Cell Biol 22: 213–224

Stallaert W, Brüggemann Y, Sabet O, Baak L, Gattiglio M, Bastiaens PIH (2018) Contact inhibitory Eph signaling suppresses EGF-promoted cell migration by decoupling EGFR activity from vesicular recycling. Sci Signal 11

Tanigawa G, Orci L, Amherdt M, Ravazzola M, Helms JB, Rothman JE (1993) Hydrolysis of bound GTP by ARF protein triggers uncoating of Golgi-derived COP-coated vesicles. J Cell Biol 123: 1365–1371

Taylor RJ, Carrington J, Gerlach LR, Taylor KL, Richters KE, Dent EW (2020) Double UP: A Dual Color, Internally Controlled Platform for. Front Mol Neurosci 13: 82

Tie HC, Ludwig A, Sandin S, Lu L (2018) The spatial separation of processing and transport functions to the interior and periphery of the Golgi stack. Elife 7

Tojima T, Miyashiro D, Kosugi Y, Nakano A (2023) Super-Resolution Live Imaging of Cargo Traffic Through the Golgi Apparatus in Mammalian Cells. Methods Mol Biol 2557: 127–140

Tojima T, Suda Y, Ishii M, Kurokawa K, Nakano A (2019) Spatiotemporal dissection of the *trans*-Golgi network in budding yeast. J Cell Sci 132

Wang H, Vilela M, Winkler A, Tarnawski M, Schlichting I, Yumerefendi H, Kuhlman B, Liu R, Danuser G, Hahn KM (2016) LOVTRAP: an optogenetic system for photoinduced protein dissociation. Nat Methods 13: 755–758

Weigel AV, Chang CL, Shtengel G, Xu CS, Hoffman DP, Freeman M, Iyer N, Aaron J, Khuon S, Bogovic J et al (2021) ER-to-Golgi protein delivery through an interwoven, tubular network extending from ER. Cell 184: 2412–2429.e2416

Weller SG, Capitani M, Cao H, Micaroni M, Luini A, Sallese M, McNiven MA (2010) Src kinase regulates the integrity and function of the Golgi apparatus via activation of dynamin 2. Proc Natl Acad Sci U S A 107: 5863–5868

Wong-Dilworth L, Rodilla-Ramirez C, Fox E, Restel SD, Stockhammer A, Adarska P, Bottanelli F (2023) STED imaging of endogenously tagged ARF GTPases reveals their distinct nanoscale localizations. J Cell Biol 222

Zink S, Wenzel D, Wurm CA, Schmitt HD (2009) A link between ER tethering and COP-I vesicle uncoating. Dev Cell 17: 403–416

