## Supplementary figures and images for "RudLOV—a new optically synchronized cargo transport method reveals unexpected effect of dynasore"

### Figure S1.jpeg

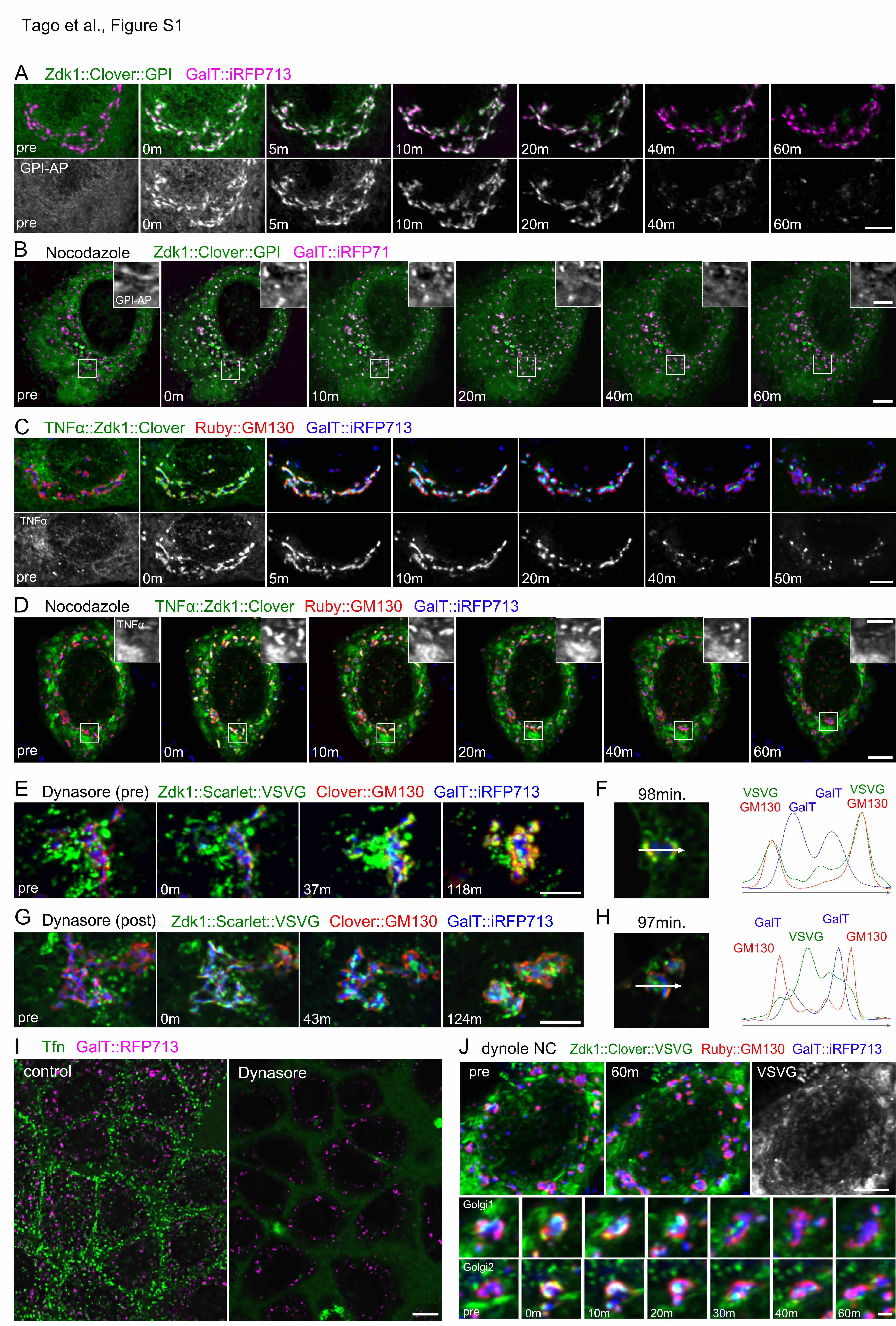
